# Modelling Signalling Networks from Perturbation Data

**DOI:** 10.1101/243600

**Authors:** Mathurin Dorel, Bertram Klinger, Anja Sieber, Anirudh Prahallad, Torsten Gross, Evert Bosdriesz, Lodewyk Wessels, Nils Blüthgen

## Abstract

**Motivation:** Intracellular signalling is realized by complex signalling networks which are almost impossible to understand without network models, especially if feedbacks are involved. Modular Response Analysis (MRA) is a convenient modelling method to study signalling networks in various contexts.

**Results:** We developed a derivative of MRA that is suited to model signalling networks from incomplete perturbation schemes and multi-perturbation data. We applied the method to study the effect of SHP2, a protein that has been implicated in resistance to targeted therapy in colon cancer, using data from a knock out and parental colon cancer cell line. We find that SHP2 is required for MAPK signalling, whereas AKT signalling only partially depends on SHP2.

**Availability:** An R-package is available at https://github.com/MathurinD/STASNet

## Introduction

Cells constantly receive external cues that are integrated by signalling networks in the cell to direct their cell fates. The topology of those signalling networks is understood to a great extend (Caron *et al*., 2010, Oda *et al*., 2005). However, the complexity of these networks makes it difficult to predict what the outcome of a perturbation would be, as feedbacks and cross-talk render intuitive reasoning impossible.

During the last years, several approaches have been developed to use computational models to tackle this problem. These approaches range from Boolean models that use logical rules to abstract the interactions between the elements of the network (Grieco *et al*., 2013) to complex models based on differential equations that model details of the reaction kinetics (Raue *et al*., 2014) or more phenomenological stimulus-response kinetics (Korkut *et al*., 2015). Boolean approaches have proven useful to predict the outcome in response to major alterations such as mutations or copy number alterations, but they fail to explain more subtle differences between cells and have problems to describe dynamic processes or effects of feedbacks (Saadatpour and Albert, 2013). On the other side of the spectrum, differential equations can be used to describe the system in details. However, fitting those models requires a tremendous amount of data, limiting them to a very small scale. Intermediate approaches typically require a limited amount of data and model quantitative responses of the signalling networks to perturbations. Those methods have the major advantage of providing a way of simulating complex networks with relatively little data. Modular Response Analysis (MRA) is an example of such approaches, where the individual phosphorylation and dephosphorylation events of kinases and phosphatases are abstracted as influences between modules. MRA was first formulated as a matrix inversion problem, and the corresponding singular value decomposition approach has been used to study the activation of the MEK-ERK cascade by NGF and EGF (Santos *et al*., 2007). A maximum likelihood formulation has been developed to study regulatory interactions between signalling, proteins and mRNA (Stelniec-Klotz *et al*., 2012), and refined to predict drug combinations to overcome resistance mechanisms (Klinger *et al*., 2013). A more recent approach based on Bayesian networks where prior information on the parameter can be integrated was also described (Halasz *et al*., 2016).

In this article, we describe an augmented version of MRA (Kholodenko *et al*., 2002; Klinger *et al*., 2013) that is particularly suited to model and reverse engineer signalling networks using perturbation data. We illustrate the approach by modelling the role of PTPN11 (SHP2) in EGFR signalling. To this end we reverse engineer the networks of a colon cancer cell line and a PTPN11 knock out derivative, which shows that PTPN11 is required to activate MAPK signalling, but has little influence on PI3K/AKT signalling, which contradicts previous literature (Wu *et al*., 2001). Our results show how modelling perturbation data of isogenic cell lines can help to uncover the role of individual proteins in signalling networks. We provide our method as an open-source R-package called STASNet (STeady-STate Analysis of Signaling Networks).

## Materials and Methods

### Perturbation response

We consider a dynamical system for the (log-transformed) read-outs ***x***, which represent the phosphorylation status of the considered kinases,

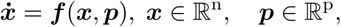

with parameters ***p*** and stable steady state ***φ***(***p***). This section describes how to infer the strength of direct interaction between different nodes (components of ***x***) from measurements of the steady states at different parameter values.

Let any type of experimental perturbation of the system be represented by a change of a specific parameter *k* from its unperturbed state 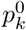 to a new value 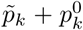. Furthermore, indicator **Δ*p*** ∈ {0,1}^p^ shall denote which parameters *p_k_* were perturbed (Δ*p_k_* = 1) or not (Δ*p_k_* = 0) in a given experimental setting. Thus, any combination of types of perturbations, represented by **Δ*p***, is associated with parameter values 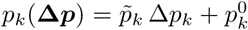. Also, we define local response and sensitivity coefficients as

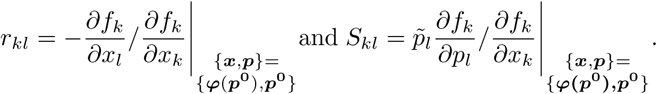

Response coefficient *r_kl_* quantifies the strength of the direct interaction from node *l* to node *k* and sensitivity coefficient *S_kl_* quantifies the effect that a perturbation on parameter *l* will have on node k. Many of those coefficients are *a priori* known to be zero, whenever direct connectivity between certain node pairs can be excluded (prior network knowledge) or perturbations are known to be specific to certain network nodes. Our goal is to determine the remaining unknown entries from experimentally measured steady state changes upon perturbation **Δ*p***, which we shall denote as ***R*** *∈* ℝ^n^. To make the connection, we realize that

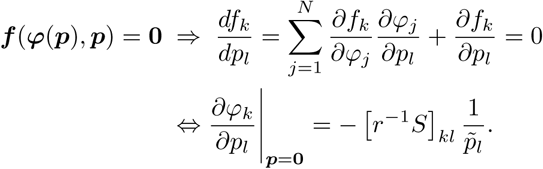

Assuming that perturbations are sufficiently small, the steady state function becomes approximately linear,

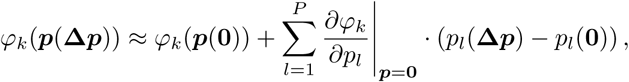

the previous equations combine to

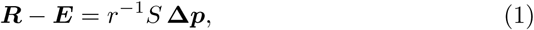

where ***E*** is an unknown error vector that compensates for measurement error and non-linearities in the steady state function ***φ***(***p***).

We analyse a perturbation experiment comprised of stimulations or inhibitions, each of which acts specifically on a single reactant. We can thus simplify the above equation by defining **Δ*p*** such that the first s entries correspond to the possible stimulations and the last i entries to the possible inhibitions, denoted as **Δ*p****^T^* = [**Δs Δ*i***]*^T^*. Partitioning the sensitivity matrix accordingly 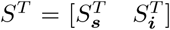, we can write *S* **Δ*p*** = *S****_s_* Δ*s*** + *S****_i_* Δ*i***. Because the perturbations are node-specific, the *k*-th row of *S****_s_*** or *S****_i_*** is either a zero vector or has a single non-zero entry, 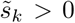 or 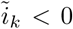 respectively. Thus, for stimulations we define ***s*** = *S****_s_* Δ*s***, where *s_k_* equals 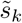 if kinase *k* was stimulated, and zero otherwise. An inhibitor however only decrease its target kinase’s signal if that kinase has a basal activity at all. So, we define ***i*** = *S****_i_* Δ***i*, where *i_k_* equals 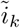 if kinase *k* was inhibited and basally active, and zero otherwise. To check for a node’s basal activity we inspect whether its exclusive inhibition provokes any measurable response. With this, Eq. (1) can be rewritten as

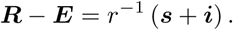

Moreover, the data indicated that kinase inhibition gives rise to a prominent non-linear behaviour that is not yet included in the above equation. In addition to the reduction of the innate signal send out by the targeted molecule, an effect which is conveyed by vector ***i***, an inhibition also reduces the ability of the inhibited node to relay upstream signals to downstream nodes. To take this second effect into consideration, we downscale the strengths of all outgoing links by factor exp(*i_l_*), 0 < exp(*i_l_*) ≤ 1 and call the altered response coefficients 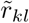, with 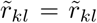 exp(*i_l_*) ∀*k ≠ l*, and 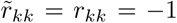, and adapt the earlier equation

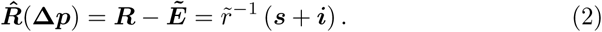

Having corrected for this most important non-linearity, we now assume that 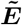 entries represent measurement error alone and consider them as *iid*-samples of a Gaussian distribution.

### Parameter estimation

The experimental data represents the measurements of the steady state differences due to different types of perturbations, **Δ*p***^1^… **Δ*p***^q^, which we combine in matrix R^expt^ ∈ ℝ^n×q^. Accordingly, we combine the expressions in Eq. (2) to form 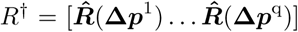, which is a symbolic representation of the perturbation response as a function of the system’s parameters *R*^†^ = *R*^†^(*r,* ***s, i***). Since the error terms in Eq. (2) are assumed to be normally distributed, we can thus fit the parameters by minimizing the sum of weighted squared residuals

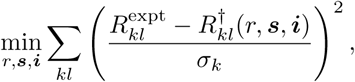

where *σ_k_* represents the analyte-specific standard deviation estimated from the replicate measurements of the data. This is done with the Levenberg-Marquard algorithm (levmar v.2.5 Lourakis, 2004). To ensure convergence to the global optimum, the procedure is repeated over various starting parameters generated through improved Latin Hypercube Sampling, as implemented in the R package Ihs.

Note, that this formulation allows to directly include prior-knowledge about the variables, by setting them to their respective values (typically zero). Furthermore, it can handle missing data by simply leaving out all those entries where 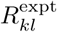 is not known.

### Parameter sensitivity analysis

STASNet implements profile likelihood as described in Raue *et al*., 2009. Briefly, for each parameter a likelihood profile is generated by refitting the model with one parameter kept constant to a series of values around its optimum. These profiles are used to detect nonidentifiable parameters, which correspond to an over-parametrisation of the model relative to the data available and to asses the reliability of parametrisation by calculating the pointwise confidence interval.

### STASNet implementation and availability

The core functions of STASNet are implemented in C++ which are accessible via wrapper functions in R. STASNet is available as an R package under https://github.com/MathurinD/STASNet.

### Generation of perturbation data input

All cell lines are derived from the colon cancer cell line Widr. The cell lines were cultured in RPMI Medium 1640 (Gibco, Life Technologies) with indicator, L-glutamine 20nM, 100U/ml Penicillin and Streptomycin, and 10% FCS. After serum starving the cells for 24 h the cells were treated 90 minutes before harvesting with an inhibitor (MEK: AZD6244 1*μ*M, PI3K: GDC0941 1*μ*M, pan-RAFi: Sorafenib 10*μ*M, BRAF600E: Vemurafenib 10*μ*M; all SelleckChem) or DMSO, and 30 minutes before harvesting with a ligand (EGF 20 ng/mL, NRG1 25 ng/mL, HGF 25 ng/mL; all R&D System) or BSA. The cells were then lysed with Bio-Plex Pro Cell Signaling Reagent Kit (Bio-Rad) and multiplexed by incubating with antibody-coated magnetic beads as described previously (Klinger *et al*., 2013) detecting the following signals: AKT^S473^, ERK1/2^T202,Y204/T185^’^Y187^, MEK1^S217^’^S221^, p90RSK^S380^, GSK3A/B^S21/S9^, RPS6^S235/S236^ and mTOR^S2448^. The plates were read using Bio-Plex Protein Array system (Bio-Rad, Hercules, CA) and the resulting .lxb files were processed using the R package lxb and a custom script to generate MIDAS-formatted files (as used by DATARAIL Saez-Rodriguez *et al*., 2008) (Supplementary S2).

## Results

### A pipeline to model signal transduction networks from perturbation data

We developed a computational pipeline to model signalling networks from perturbation data using Modular Response Analysis. It is based on a previously established maximum-likelihood formulation of MRA (Stelniec-Klotz *et al*., 2012), and extensions that cover the effects of inhibitors on interactions and a reparameterization to account for non-identifiable parameters (Klinger *et al*., 2013), for details confer Materials and Methods.

In the following we describe the typical steps to model and analyse the data (see also Fig. 1): From a prior knowledge network of the signalling pathways and the experimental layout (i.e. which signalling nodes are stimulated or inhibited, and which nodes are measured), our algorithm constructs an MRA-based model. Briefly, the algorithm constructs a symbolic local response matrix (i.e. a normalised Jacobian matrix), inverts this matrix, and then computes symbolic expressions for the global response matrix. Next, data on signalling node activity (such as phosphorylation of kinases) before and after perturbation by e.g. ligands and inhibitors are used to estimate the parameters using a maximum-likelihood approach. By iteratively probing if addition of links significantly increases fit quality, or removal of links does not alter fit quality significantly, the network can be refined. Once a model with reasonable fit quality is generated, its parameters can be analysed (and confidence intervals computed) using profile likelihood. Models that were generated for different cell lines can be compared, and model simulations can be used to predict unseen perturbations.

**Figure 1:**
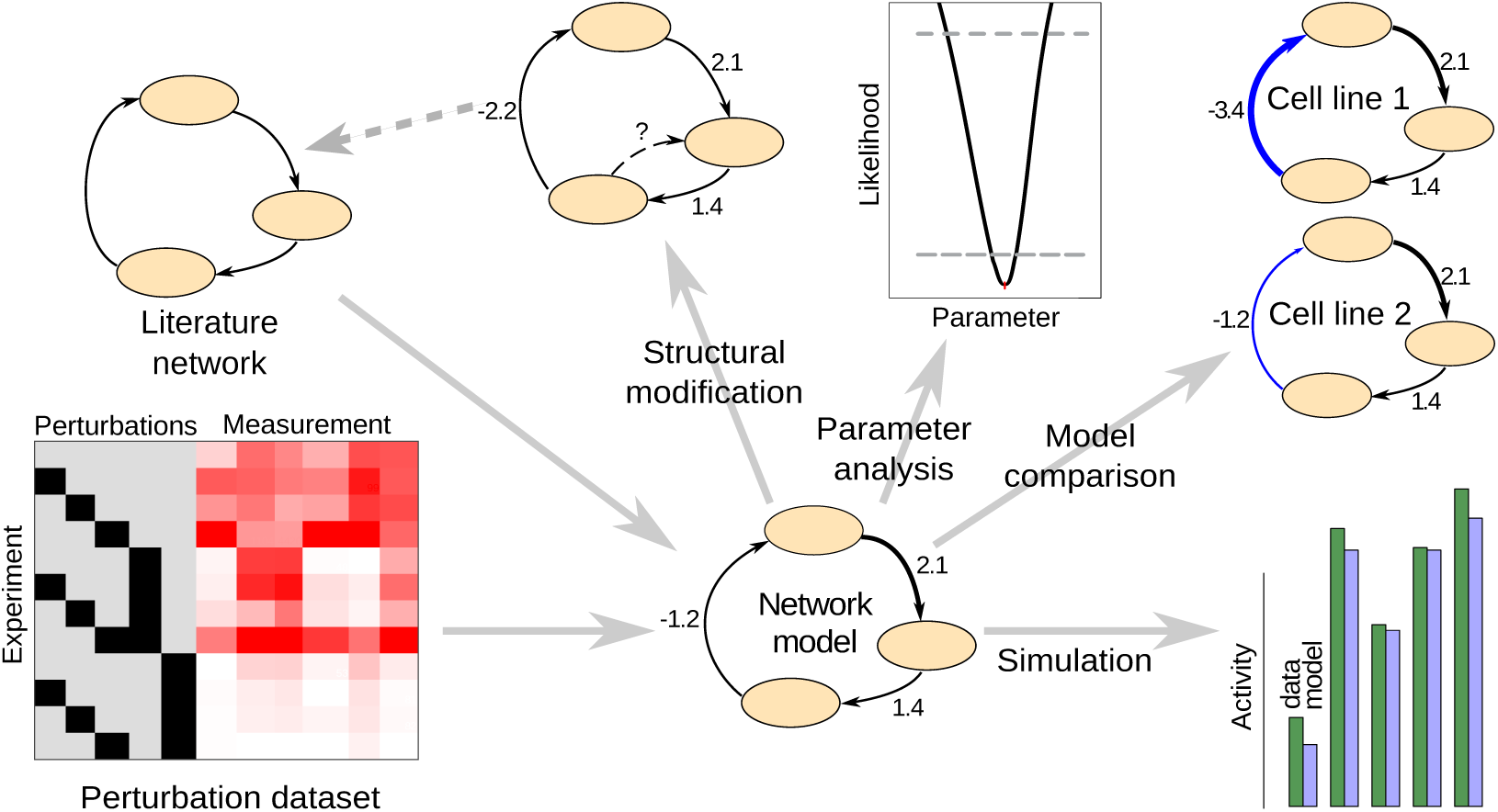
Workflow of the STASNet package. STASNet uses network structure, experimental design and perturbation data as inputs to generate a signalling network model. This model can then be used to suggest modifications in the network structure that are necessary to explain the data (Structural modification), be analysed with profile likelihood (Raue *et al*., 2009) (Parameter analysis), compared to models from other cells (Model comparison) or used to simulate the response of the network to - also novel - perturbations (Simulation).

### An *in-silico* example

To illustrate the application of STASNet, we applied it to *in-silico* perturbation data, where we simulated perturbations in a model of a signalling network using ordinary differential equations. To generate data, we used a network model to simulate a cascade of three nodes (A, B, C), where the last node (C) inhibits the first (A), and a “ligand” S stimulates A. The model also included an inhibitor of A (Figure 2A). We simulated four conditions: control conditions, stimulation of A by S, inhibition of A by *i*_A_, and a combination of the two perturbations. Our STASNet pipeline requires three input files (Figure 2C) containing 1) the perturbation data in MIDAS format (.csv) (described in Saez-Rodriguez *et al*., 2008), 2) the network structure as a two-columned table describing the links (.tab), 3) the nodes with basal activity (.dat). The latter file lists all signalling nodes in the network that have basal activity, i.e. those nodes where inhibition alone leads to decreased activity in downstream nodes, as discussed below.

**Figure 2:**
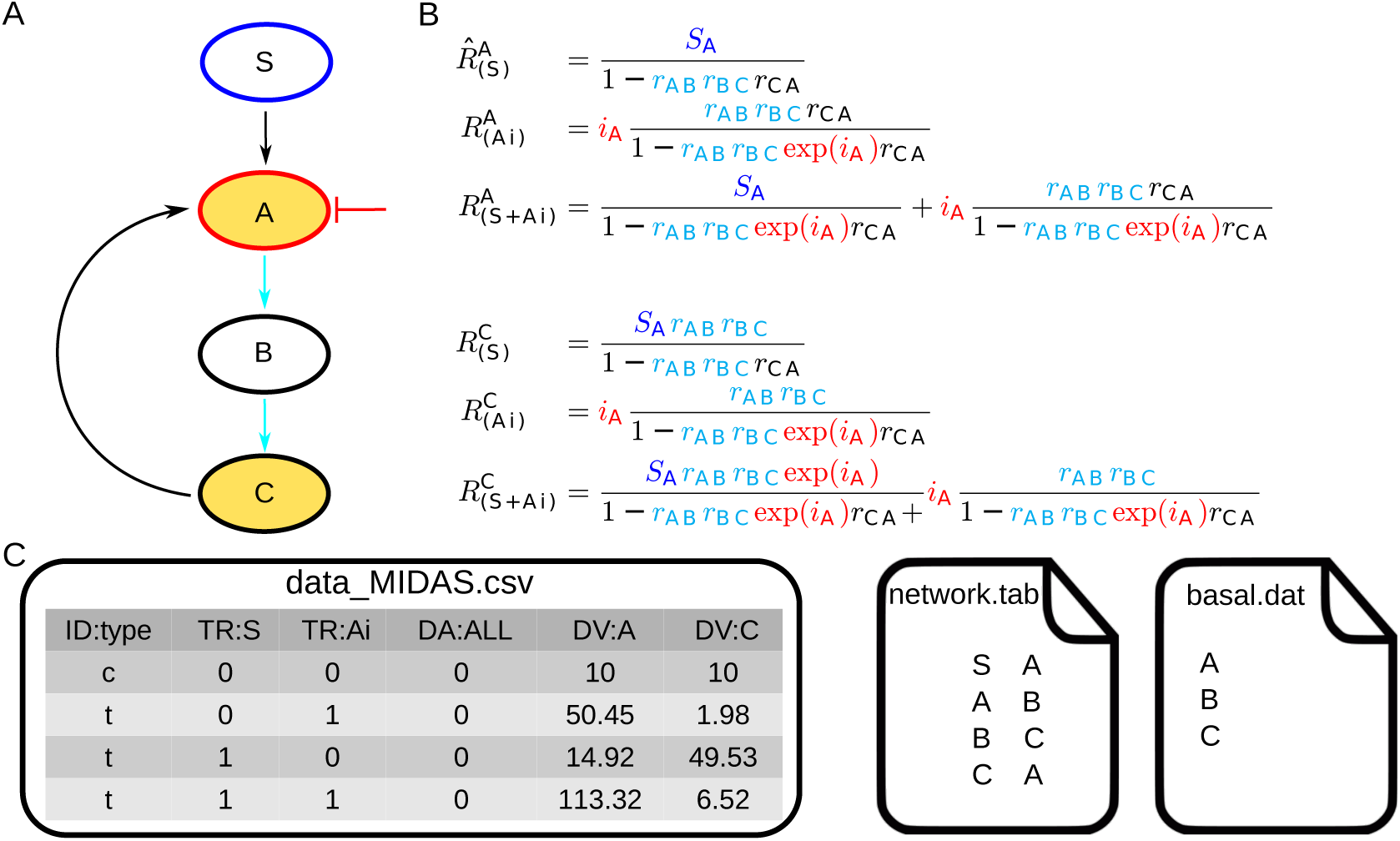
Minimal STASNet example. Minimal example of package inputs and the underlying symbolic equations. **A** Example of a perturbation scheme. **B** Symbolic equations of selected global responses generated by the package with structurally identifiable parameters colorized. As B is neither measured nor perturbed, *r_AB_ r_BC_* is reduced to one parameter by STASNet. The inhibitory parameters are depicted in red. **C** Input files for STASNet: the data_MIDAS.csv file specifies the experimental layout in (A) (stimulation of S, inhibition of A, and measurement of A and C), network.tab contains the network structure and basal.dat the nodes that have basal activity (see Material and Methods).

From these input files, STASNet derives the symbolic local response matrix, from which, after inversion, it computes the global response coefficients symbolically which are then used to model the data. These response coefficients represent the log-fold-changes for each measured node in response to the perturbations.

Examples of these response coefficients for our *in silica* example are shown in Figure 2B. The first equation describes the response of the network to stimulation S. It should be noted that the two local response coefficients *r_AB_* and *r_BC_*, that describe the strength of signalling between A and B, and B and C, respectively, cannot be inferred from the data separately, as node B is not measured. STASNet analyzes the equations to re-parametrize the model using Gaussian elimination, as described previously (Klinger *et al*., 2013). In brief, the algorithm detects parameters that only occur together in products or ratios, and defines new identifiable parameter combinations. In our example, the re-parametrization defines a new parameter that represents the product *r_AB_* · *r_BC_*.

The second equation in Figure 2B shows the response to the inhibition of A. Here we assume that A has basal activity, i.e. the inhibitor perturbs downstream signalling with a parameter 1*_A_* independently of upstream activation of A. Without basal activity of A, this response would be 0. In addition to being a perturbation on downstream nodes, inhibition of A also alters the local response coefficients, which we model by multiplying the respective control coefficients with exp(*i_A_*). This altered response coefficient is visible in the effect of feedbacks (subtrahends in the denominator).

### Perturbation data for RTK signalling in a colorectal carcinoma cell line

Next, this pipeline can be used to study specific questions on signalling. We decided to study the role of PTPN11, a phosphatase that is important in receptor tyrosine (RTK) signalling, and has been implicated in feedback control of EGFR signalling and drug resistance (Prahallad *et al*., 2015). To do so, we chose to generate a model of RTK signalling in a colorectal cancer cell line Widr containing an activating BRAF^V600E^ mutation, and then compare this model with a model of the same cell line, where PTPN11 is inactivated using CRISPR/Cas9. To parametrize the model, we measured the phosphorylation state of seven kinases involved in MAPK and PI3K signalling, and their response to three ligands (EGF, HGF and NRG1) and two inhibitors (MEK and PI3K inhibitors), alone and in combination (see Figure 2A). The experimental design was such that we serum-starved the cells, incubated them with inhibitors or their solvent control for one hour, and subsequently stimulated them with the ligands or their solvent. Half an hour later, cells were lysed and phospho-proteins were measured using bead-based ELISAs.

### Adapting the literature signalling network

Apart from the perturbation data, our modelling framework requires the signalling network topology as input. The MAPK and PI3K pathways are well studied, which allowed us to generate a literature-derived interaction network (Fig 3B). The network consists of the three ligands, their receptors, RAS, PI3K and RAF as unmeasured signalling intermediates, the measured pathway components of AKT and MAPK signalling and pathway targets that include mTOR (as an AKT target) and GSK3, p90RSK and RPS6 as common targets of both signalling pathways. In addition, we included two well-studied feedback loops in MAPK signalling (ERK→RAF and ERK→EGFR) that are known to play a role in drug resistance (Fritsche-Guenther *et al*., 2011,Klinger *et al*., 2013).

**Figure 3:**
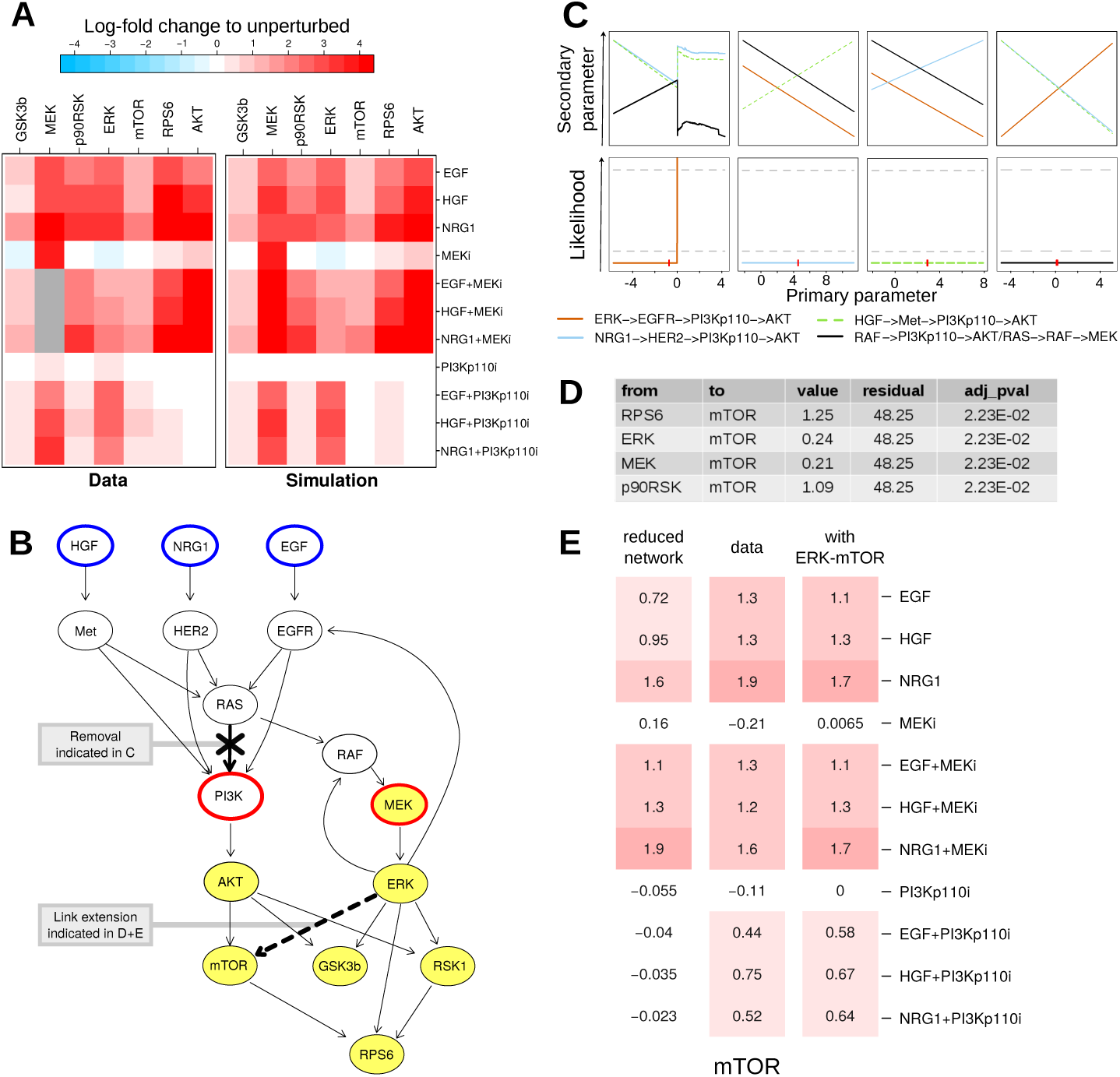
Building a model using STASNet. **A** Experimental data of human CRC Widr cell line and simulation results from the initial literature-derived network (depicted in (B)). Grey squares indicate missing values. **B** Kinase interaction network, including experimental design with measured (yellow), stimulated (blue) and inhibited (red) nodes. Updates from the initial network are indicated by bold links: removal RAS→PI3K link, and extension of ERK→mTOR link. **C** Profile likelihood of paths containing a link between receptors and AKT computed for the initial network. Red marks indicate the fitted value of the primary parameter. **D** Top links suggestion with equal improvement from the link extension feature. **E** Comparison of the mTOR response simulations with or without the ERK→mTOR link to the data.

The model contained 19 parameters that represent either entries of the local response matrix (or lumped combinations of them), or inhibitor strengths. When estimating these 19 parameters using a maximum likelihood procedure, the weighted sum of squared residuals was 52, which is compatible with the 74 measurements. An interesting aspect of the network are the different modes of activation of the two pathways: while the MAPK pathway is solely activated through RAS, AKT is activated both in a RAS-independent and -dependent way (Hemmings and Restuccia, 2012). However, as we neither measure or perturb elements of these two pathways leading to AKT activation, their parameter values cannot be estimated independently. This is known as structural non-identifiability. Our pipeline allows to calculate the profile likelihood for the model, which shows the change in maximum likelihood when one (primary) parameter is varied (Raue *et al*., 2009). When plotting the profile likelihoods and the optimized parameters, this structural non-identifiability is directly visible by flat profile likelihoods and compensatory changes in related (secondary) parameters (Fig 3C). To resolve this, our pipeline allows to sequentially remove links from the model by re-fitting all possible models with one link removed and comparing the resulting log-likelihoods. In our example, the model can be reduced by removing one of the links HER2→PI3K, RAS→PI3K, Met→PI3K or EGFR→PI3K without changes in the likelihood (Supplementary S1). As all three models have the same likelihood, we decided to remove the link RAS→PI3K as it separates the PI3K and RAF cascades (Fig 3D) and allows for more straight forward interpretation of the parameters; it also leads to a numerically more stable model. This results in a network model where all links are identifiable.

When comparing the data and the model fit, we noticed that although in general the data can be reproduced reasonably well, there are some discrepancies for mTOR upon PIK3 inhibition (see Fig 3E reduced network vs. data). To investigate if any additional links can resolve these discrepancies, we use the extension suggestion feature of STASNet (Supplementary S1). Briefly, this features tests all possible links, ranks them according to their likelihood and evaluates their significance. We noticed that adding any of the links RPS6→mTOR, ERK→mTOR, MEK→mTOR, or p90RSK→mTOR does not improve fit quality (Fig 3C). Since the experimental setup did not permit us to discriminate these links we searched the literature and found that only of ERK an activation of mTOR by inhibiting mTOR inhibitory complex protein TSC2 via phosphorylation at serine 664 is described (Rolfe *et al*., 2005). We thus updated the network to include this link, which led to an improved fit (Fig 3E with ERK→mTOR). The final model is displayed in Fig. 3B.

### Analysis of a SHP2 knock-out with STASNet

After having established a model for the cell line with a wild-type SHP2, we next aimed to model the network when SHP2 is inactivated. For this, we used a SHP2 KO cell line derived from the Widr cell line using CRISPR/Cas9 (Prahallad *et al*., 2015). SHP2 (PTPN11) is a phosphatase that binds to the tyrosine kinase receptors through adaptors such as GAB1 and participates in the activation of the MAPK cascade by relieving inhibitions on RAS and RAF. SHP2 has been shown to be reactivated in BRAF inhibition resistant cell lines (Prahallad *et al*., 2015) (Fig 4A).

**Figure 4:**
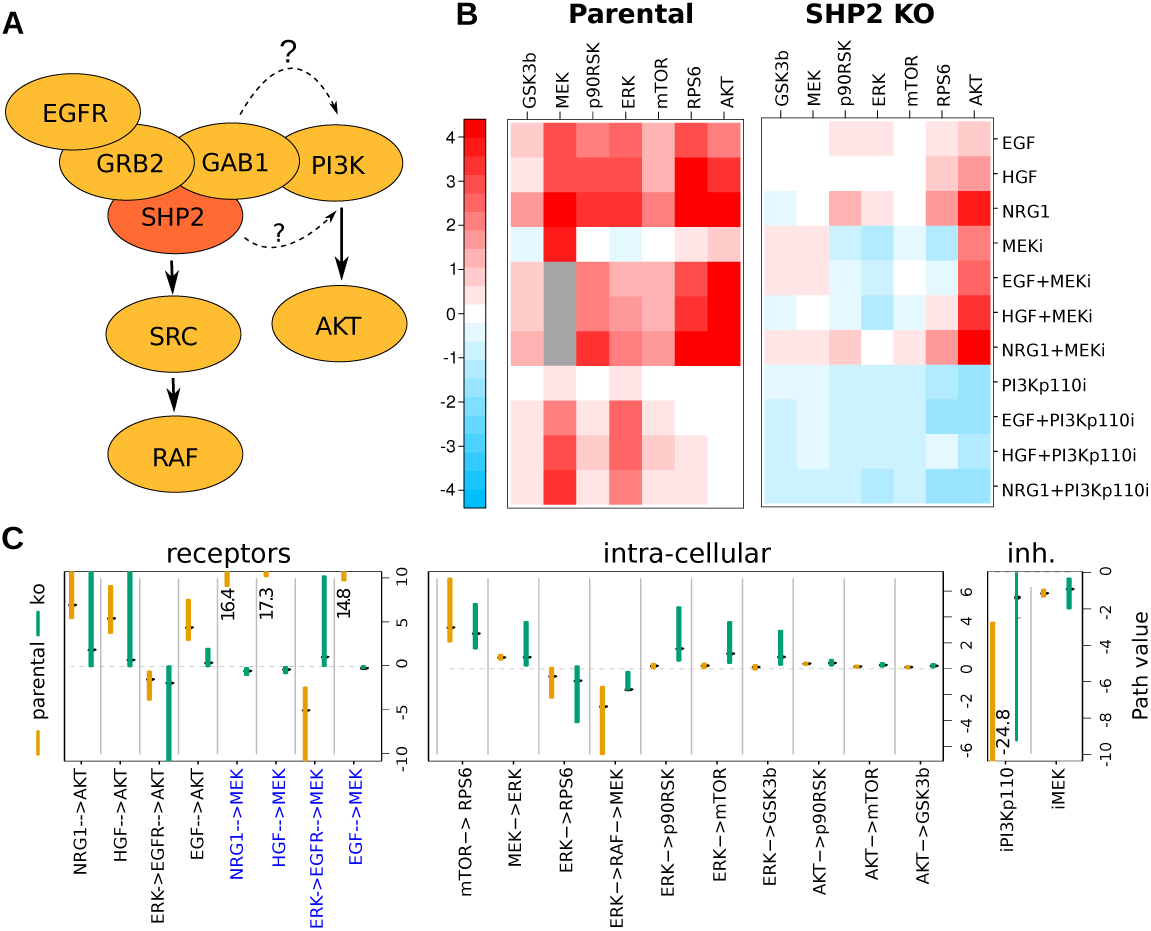
Analysis of a SHP2 KO with STASNet. **A** Literature knowledge of the role of SHP2 in EGFR signalling **B** Perturbation data for the parental and SHP2 KO cell lines **C** Parameters with pointwise confidence interval of the respective models. All links from receptor to MEK are affected by SHP2 KO (dark blue).

We applyied the same perturbation experiments to the SHP2 KO cells and compared the response to the parental cell line (see Fig 4B). It is evident that the knock out led to a reduction of phosphorylation response throughout the network. We then parametrized a model for signalling in the SHP2 KO cell line, using the network derived for the parental cell line, and obtained a maximum likelihood fit sufficiently complying with the experimental error. We then quantitatively compared the parameters fitted for the SHP2 KO to those fitted for the parental cell line (Fig 4C) using confidence intervals for the parameters obtained by profile likelihood. For intra-cellular paths and inhibitors the values between parental and KO cell lines did not significantly differ (Fig 4C middle and lower panel). For the receptor connecting paths we found several parameters that were significantly (and qualitatively) different between the two cell lines. Four of these parameters correspond to all paths in the network that connect the receptors to MAPK signalling. While these parameters are numerically large in the parental cell line, they are close to zero in the SHP2 KO (Fig 4C, upper panel indicated in blue). This confirms the known role of SHP2 as being between the receptor and the activation of RAF (Fig. 4A). Accordingly, the ERK→RAF→MEK feedback is still functional in the SHP2 KO. Surprisingly however, the parameters corresponding to the activation of AKT by two of the three ligands are not significantly altered, suggesting that SHP2 is only partly required for the activation of the PI3K/AKT pathway (Fig. 4C). Interestingly, the ERK→EGFR→AKT crosstalk is also still functional which indicates that ERK regulation of EGFR does not depend on SHP2.

### Prediction of the impact of RAF inhibition

Having cell-specific models generated for the parental and SHP2 KO, we can then ask how other perturbations would affect the networks. RAF is an important therapeutic target for which two main classes of inhibitors exist. Some inhibitors, such as Vemurafenib, target specifically mutant BRAF (BRAF^V600E^, Bollag *et al*., 2010), whereas others, such as Sorafenib, are pan-RAF inhibitors that inhibit all RAF isoforms (Wilhelm *et al*., 2004). The Widr cell system that we study harbours the BRAF^V600E^ so we could investigate the effects of these two inhibitor classes in our model and experimentally validate them afterwards.

As we calibrated our model on data where RAF activity was neither measured nor perturbed, the RAF**→**MEK link is not directly fitted but in parameter combinations. We therefore had to augment our model, by including a new node, BRAF^V600E^ mutation that is connected to MEK and receives no upstream signal (Fig. 5A), as the BRAF^V600E^ mutation renders BRAF insensitive to upstream and feedback signals (Friday *et al*., 2008). Moreover since we had to give an reasonable but arbitrary parameter value for the inhibition strength for both RAF inhibitors (set to 3), the resulting predictions can only be understood qualitatively.

**Figure 5:**
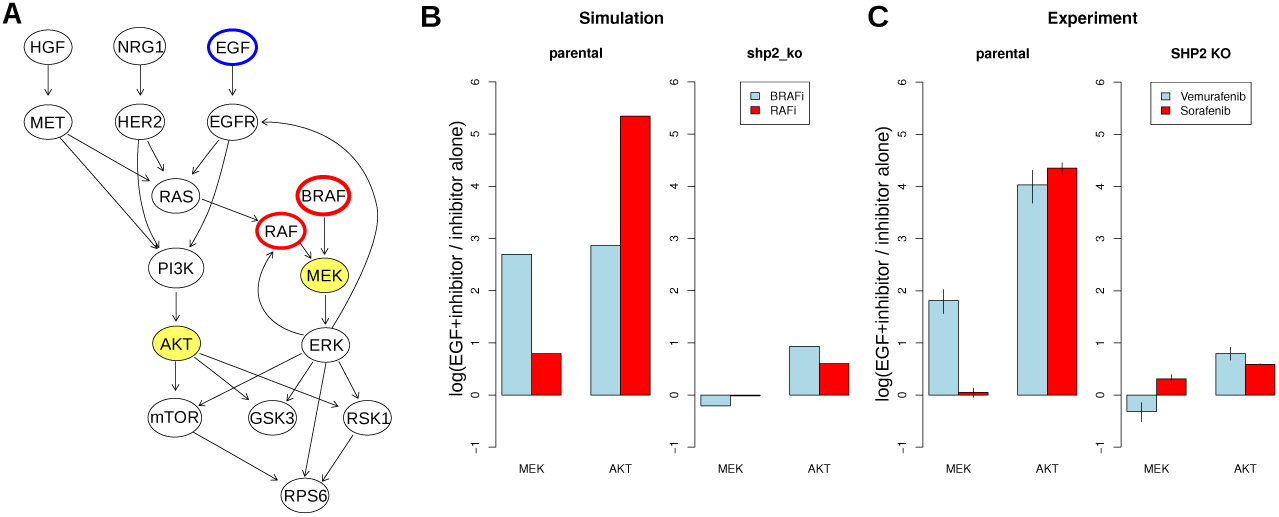
Prediction of RAF inhibitors effects. **A** Localisation of RAF and BRAF inhibition in the network derived for the Widr cell line; measured/predicted nodes are highlighted in yellow. **B** STASNet simulation of the impact of BRAF^V600E^ and pan-RAF inhibitors on both cell lines **C** Experimental measurements of the impact of Vemurafenib (BRAF^V600E^ inhibitor) and Sorafenib (pan-RAF inhibitor) on the parental and SHP2 KO cell lines (error bars in S.D., n=2).

One of the resistance mechanisms for RAF inhibition is the reactivation of the RAF-MEK-ERK signalling pathway and the activation of AKT by loss of feedback inhibition of the EGF receptor (Prahallad *et al*., 2012, Klinger *et al*., 2013). We therefore decided to simulate to what extend EGF stimulation rescues MEK and AKT phosphorylation upon RAF inhibition. In our simulation, we noticed qualitative differences between the two inhibitors in our wild-type cell lines (Fig. 5B). While EGF stimulation hyperactivates AKT for both inhibitors, it only rescues MEK phosphorylation after treatment with the BRAF^V600E^ specific inhibitor. This is consistent with the idea that this inhibitor specifically blocks the mutant allele, while the non-mutated allele and other isoforms can still relay the signal. When comparing the simulations of the two different cell line models it can be noted that in the parental cell line, both MEK and AKT are upregulated with EGF, whereas in SHP2 KO this upregulation is completely blocked for MEK, and is attenuated for AKT (Fig. 5B).

To confirm our model prediction, we performed experiments in which we preincubated the cells with either Vemurafenib (BRAF^V600E^ inhibitor) or Sorafenib (pan-RAF inhibitor) for 90 min, and then stimulated cells with EGF for 30 min, and measured AKT and MEK phosphorylation (Fig. 5C). These data are in qualitative agreement with our model predictions, confirming the disruption of the EGF–¿MEK path and the attenuation of the EGF–¿AKT path in the SHP2 KO.

## Discussion

Perturbation-response data sets are key for the analyses of signalling networks, and many different approaches have been developed to generate computational models from such systematic data sets. Most of these approaches binarize data and use logic approaches to describe the data (Morris *et al*., 2010), or alternatively they use dynamic information to fit quantitative data (Raue *et al*., 2014). In this article, we describe an approach that uses an augmented version of MRA (Kholodenko *et al*., 2002; Klinger *et al*., 2013), that allows the generation of semi-quantitative models from snapshot perturbation-data. We developed an R package called STASNet, that implements this approach and provides analysis functions to improve and compare models.

In this article, we applied STASNet to model the effect of a SHP2 knock-out on the MAPK and PI3K signalling network. By comparing the parameters of the MRA models for the two isogenic cell lines we could recover the known role of SHP2 in mediating MAPK signalling. However, it was unclear if SHP2 is required to activate PI3K/AKT signalling in receptor tyrosine kinase/EGFR signalling. We found that PI3K/AKT signalling triggered by the receptors HER2 and HGF is largely independent of SHP2, whith EGFR being partly dependent.

SHP2 has been implicated in resistance of BRAF mutant colorectal carcinoma, where loss of feedback to SHP2/EGFR leads to reactivation of MAPK signalling after treatment with BRAF inhibitors (Prahallad *et al*., 2015). Our study confirms that SHP2 has a major role in re-activation of MAPK signalling, as our model predictions show and the data confirm that MEK phosphorylation cannot be recovered in SHP2 KO with EGF. Furthermore the model predicts and the data confirms that with functional SHP2, pan-RAF inhibitors also prevent activation of the MAPK pathway, and may be considered an alternative treatment option to prevent resistance.

The signalling focus of our method is set on the intermediate early time points when phosphorylation events have reached a quasi steady-state and transcriptional feedbacks are not yet appearing. We think that this intermediate state is useful to predict the cell response to drugs directed on the signalling network components and to identify potential resistance mechanisms. MRA has also already been used to compare different cell lines to explain drug resistance and predict ways to overcome them (Klinger *et al*., 2013). STASNet provides a convenient way of applying such methods on various perturbation data.

Few other programs exist to deal with single time point steady state data. CellNOptR (Terfve *et al*., 2012) can be applied to such data after discretisation or normalisation of the data, which implies to define thresholds and might require external data. Other more precise approaches like CNORode (Terfve *et al*., 2012) or Data2Dynamics (Raue *et al*., 2014) rely on ordinary differential equations and require more data to be parametrized, limiting their application to small networks.

To conclude, STASNet provides a convenient R-package to generate MRA-models using a maximum likelihood approach for single time point signalling data. The package can be obtained at https://github.com/MathurinD/STASNet, and example data and the analysis scripts are available within the package. R scripts that can reproduce the analysis in this paper are in the supplementary information.

## Supporting Information

S1 File Knitr script with input to generate the figures 3 and 4

S2 File zip file containing lxb files (raw data) and R scripts to process them

S3 File zip file containing the data and script used for the RAF inhibitors simulation, and the lxb files for the confirmation experiment

## References

Bollag, G., Hirth, P., Tsai, J., Zhang, J., Ibrahim, P. N., Cho, H., Spevak, W., Zhang, C., Zhang, Y., Habets, G., Burton, E. A., Wong, B., Tsang, G., West, B. L., Powell, B., Shellooe, R., Marimuthu, A., Nguyen, H., Zhang, K. Y. J., Artis, D. R., Schlessinger, J., Su, F., Higgins, B., Iyer, R., D’Andrea, K., Koehler, A., Stumm, M., Lin, P. S., Lee, R. J., Grippo, J., Puzanov, I., Kim, K. B., Ribas, A., McArthur, G. A., Sosman, J. A., Chapman, P. B., Flaherty, K. T., Xu, X., Nathanson, K. L., and Nolop, K. (2010). Clinical efficacy of a RAF inhibitor needs broad target blockade in BRAF-mutant melanoma. Nature, 467(7315), 596–9.

Caron, E., Ghosh, S., Matsuoka, Y., Ashton-Beaucage, D., Therrien, M., Lemieux, S., Perreault, C., Roux, P. P., and Kitano, H. (2010). A comprehensive map of the mTOR signaling network. Molecular systems biology, 6, 453.

Friday, B. B., Yu, C., Dy, G. K., Smith, P. D., Wang, L., Thibodeau, S. N., and Adjei, A. A. (2008). BRAF V600E disrupts AZD6244-induced abrogation of negative feedback pathways between extracellular signal-regulated kinase and Raf proteins. Cancer Research, 68(15), 6145–6153.

Fritsche-Guenther, R., Witzel, F., Sieber, A., Herr, R., Schmidt, N., Braun, S., Brummer, T., Sers, C., and Blüthgen, N. (2011). Strong negative feedback from Erk to Raf confers robustness to MAPK signalling. Molecular systems biology, 7(489), 489.

Grieco, L., Calzone, L., Bernard-Pierrot, I., Radvanyi, F., Kahn-Perlès, B., and Thieffry, D. (2013). Integrative Modelling of the Influence of MAPK Network on Cancer Cell Fate Decision. PLoS Computational Biology, 9(10), e1003286.

Halasz, M., Kholodenko, B. N., Kolch, W., and Santra, T. (2016). Integrating network reconstruction with mechanistic modelling to predict cancer therapy. Science Signaling, 9(455).

Hemmings, B. A., and Restuccia, D. F. (2012). PI3K-PKB/Akt pathway. Cold Spring Harbor Perspectives in Biology, 4(9).

Kholodenko, B. N., Kiyatkin, A., Bruggeman, F. J., Sontag, E., Westerhoff, H. V., and Hoek, J. B. (2002). Untangling the wires: a strategy to trace functional interactions in signaling and gene networks. Proceedings of the National Academy of Sciences of the United States of America, 99(20), 12841–6.

Klinger, B., Sieber, A., Fritsche-Guenther, R., Witzel, F., Berry, L., Schumacher, D., Yan, Y., Durek, P., Merchant, M., Schäfer, R., Sers, C., and Blüthgen, N. (2013). Network quantification of EGFR signaling unveils potential for targeted combination therapy. Molecular systems biology, 9, 673.

Korkut, A., Wang, W., Demir, E., Aksoy, B. A., Jing, X., Molinelli, E. J., Babur, O., Bemis, D. L., Onur Sumer, S., Solit, D. B., Pratilas, C. A., and Sander, C. (2015). Perturbation biology nominates upstream-downstream drug combinations in RAF inhibitor resistant melanoma cells. Elife, 4.

Lourakis, M. (Jul. 2004). levmar:Levenberg-marquardt nonlinear least squares algorithms in C/C++. [web page] http://www.ics.forth.gr/͂lourakis/levmar/.

Morris, M. K., Saez-Rodriguez, J., Sorger, P. K., and Lauffenburger, D. A. (2010). Logic-based models for the analysis of cell signaling networks.

Oda, K., Matsuoka, Y., Funahashi, A., and Kitano, H. (2005). A comprehensive pathway map of epidermal growth factor receptor signaling. Molecular systems biology, 1(1), 2005.0010.

Prahallad, A., Sun, C., Huang, S., Nicolantonio, F. D., Salazar, R., Zecchin, D., Beijersbergen, R. L., Bardelli, A., and Bernards, R. (2012). Unresponsiveness of colon cancer to BRAF(V600E) inhibition through feedback activation of EGFR. Nature, 483, 100–3.

Prahallad, A., Heynen, G. J. J. E., Germano, G., Willems, S. M., Evers, B., Vecchione, L., Gambino, V., Lieftink, C., Beijersbergen, R. L., Di Nicolantonio, F., Bardelli, A., and Bernards, R. (2015). PTPN11 Is a Central Node in Intrinsic and Acquired Resistance to Targeted Cancer Drugs. Cell Reports, 12(12), 1978–1985.

Raue, A., Kreutz, C., Maiwald, T., Bachmann, J., Schilling, M., Klingmüller, U., and Timmer, J. (2009). Structural and practical identifiability analysis of partially observed dynamical models by exploiting the profile likelihood. Bioinformatics, 25(15), 1923–1929.

Raue, A., Steiert, B., Schelker, M., Kreutz, C., Maiwald, T., Hass, H., Vanlier, J., Tänsing, C., Adlung, L., Engesser, R., Mader, W., Heinemann, T., Hasenauer, J., Schilling, M., Hofer, T., Klipp, E., Theis, F., Klingmuüller, U., Schüoberl, B., and Timmer, J. (2014). Data2Dynamics: A modeling environment tailored to parameter estimation in dynamical systems. Bioinformatics, 31(21), 3558–3560.

Rolfe, M., McLeod, L. E., Pratt, P. F., and Proud, C. G. (2005). Activation of protein synthesis in cardiomyocytes by the hypertrophic agent phenylephrine requires the activation of ERK and involves phosphorylation of tuberous sclerosis complex 2 (TSC2). The Biochemical journal, 388(Pt 3), 973–84.

Saadatpour, A. and Albert, R. (2013). Boolean modeling of biological regulatory networks: A methodology tutorial. Methods, 62(1), 3–12.

Saez-Rodriguez, J., Goldsipe, A., Muhlich, J., Alexopoulos, L. G., Millard, B., Lauffenburger, D. A., and Sorger, P. K. (2008). Flexible informatics for linking experimental data to mathematical models via DataRail. Bioinformatics, 24(6), 840–847.

Santos, S. D. M., Verveer, P. J., and Bastiaens, P. I. H. (2007). Growth factor-induced MAPK network topology shapes Erk response determining PC-12 cell fate. Nature cell biology, 9(3), 324–30.

Stelniec-Klotz, I., Legewie, S., Tchernitsa, O., Witzel, F., Klinger, B., Sers, C., Herzel, H., Blüthgen, N., and Schäfer, R. (2012). Reverse engineering a hierarchical regulatory network downstream of oncogenic KRAS. Molecular systems biology, 8, 601.

Terfve, C., Cokelaer, T., Henriques, D., MacNamara, A., Goncalves, E., Morris, M. K., van Iersel, M., Lauffenburger, D. A., and Saez-Rodriguez, J. (2012). CellNOptR: a flexible toolkit to train protein signaling networks to data using multiple logic formalisms. BMC systems biology, 6(1), 133.

Wilhelm, S. M., Carter, C., Tang, L., Wilkie, D., McNabola, A., Rong, H., Chen, C., Zhang, X., Vincent, P., Mchugh, M., Cao, Y., Shujath, J., Gawlak, S., Eveleigh, D., Rowley, B., Liu, L., Adnane, L., Lynch, M., Auclair, D., Taylor, I., Gedrich, R., Voznesensky, A., Riedl, B., Post, L. E., Bollag, G., and Trail, P. A. (2004). BAY 43-9006 Exhibits Broad Spectrum Oral Antitumor Activity and Targets the RAF / MEK / ERK Pathway and Receptor Tyrosine Kinases Involved in Tumor Progression and Angiogenesis BAY 43-9006 Exhibits Broad Spectrum Oral Antitumor Activity and Targets the Pr. Cancer Res, 64(19), 7099–7109.

Wu, C. J., O’Rourke, D. M., Feng, G. S., Johnson, G. R., Wang, Q., and Greene, M. I. (2001). The tyrosine phosphatase SHP-2 is required for mediating phosphatidylinositol 3-kinase/Akt activation by growth factors. Oncogene, 20(42), 6018–6025.

